# Compressive axial-integrated planar scanning (CAPS) microscopy for high-speed volumetric imaging of cardiac dynamics

**DOI:** 10.64898/2026.04.21.720045

**Authors:** Xinyuan Zhang, Jichen Chai, Yi Gong, Milad Almasian, Jonathan Aaron Brewer, Alireza Saberigarakani, Jianqing Jia, Alyssa Hines, Kelli J Carroll, Yifei Lou, Yichen Ding

## Abstract

Investigating cardiac dynamics, including contractile function and intracardiac flow, requires volumetric imaging capable of resolving whole-organ events at micrometer resolution and millisecond timescales. However, the limited readout bandwidth of detectors imposes fundamental trade-offs among spatial sampling, field of view, and achievable volume rates. Here we introduce compressive axial-integrated planar scanning (CAPS) microscopy, a computational imaging framework that combines rapid light-sheet scanning, detection-side axial multiplexing with model-based reconstruction to enhance detector bandwidth utilization for high-speed volumetric imaging. Using widely accessible optical sensors and components, CAPS achieves cellular-scale resolving power across heart chambers at 200 volumes per second with an effective detector pixel rate of 5.82 GHz, representing a ∼15-fold increase in spatiotemporal throughput relative to uncompressed volumetric acquisition. Coordinated high-speed encoding and computational reconstruction further mitigate rolling-shutter distortions in CMOS sensors while preserving frame rate and intrinsic optical sectioning. We demonstrate that CAPS enables beat-resolved imaging of single-cell cardiomyocyte kinematics, chamber-scale contractile dynamics, and intracardiac hemodynamics in zebrafish larvae under both healthy and pharmacologically perturbed conditions. Collectively, these advances establish CAPS as a powerful framework for quantitative, *in vivo* characterization of coordinated and disrupted cardiac dynamics at cellular resolution, supporting high-speed volumetric interrogation of organ-level function and disease progression.

## Introduction

Volumetric imaging and quantitative analysis at micrometer resolution and millisecond timescales are critical for advancing our understanding of physiological processes and the mechanisms underlying disease pathogenesis, particularly for rapid organ-level dynamics such as cardiac contractile function and intracardiac flow^1–4^. Capturing these 4D (3D + time) *in vivo* events requires high spatiotemporal resolution across a wide field of view, demanding the transmission of millions to billions of pixels per second. Most established optical-sectioning imaging modalities acquire volumetric datasets through fully sequential sampling^5^. Although engineered illumination^6^, rapid beam steering^7,8^, multiplane and/or multifocus recording^9^, and retrospective^1,10,11^ or prospective^12^ gating have advanced optical-sectioning volumetric imaging, including for 4D cardiac dynamics, these strategies still rely on complete sampling in both space and time. Consequently, the readout bandwidth of digital detectors, defined as the maximum pixel sampling rate per second^13^, can become a major constraint, such that increasing volumetric imaging rate often reduces the achievable field of view or spatial resolution^8,14^.

In parallel, alternative strategies have sought to recover volumetric information from fewer measurements through optical encoding and computational reconstruction, in some cases leveraging compressed-sensing frameworks that exploit signal sparsity or compressibility to alleviate detector-bandwidth limitations. These efforts have proceeded primarily through either illumination-side or detection-side encoding^9,15,16^. Spatially modulated illumination approaches^17,18^, sometimes combined with synchronized axial or focal-plane sweeping^19–21^, have been explored, but many implementations remain constrained by multiple illuminations of the same volume and/or relatively slow mechanical scanning, resulting in effective volumetric acquisition windows on the order of hundreds of milliseconds to seconds. Detection-side strategies instead encode depth information in the emitted fluorescence. Engineered point-spread functions (PSFs) can encode axial information in the detected emission pattern, enabling single-shot volumetric reconstruction with computational decoding. However, many established implementations rely on task-specific PSF designs^22,23^, while more general universal-PSF formulations^24^ have thus far been demonstrated mainly numerically under linear, spatially incoherent assumptions. Other detection-side approaches include emerging light-field microscopy^25,26^, which reconstructs volumes by trading spatial sampling for angular information, but its widefield multiplexing introduces depth mixing and can reduce optical sectioning performance. Detection-side snapshot systems such as coded-aperture and lensless platforms can multiplex temporal^27^, spectral^28^, or depth^29^ information, although depth-encoding variants remain less established in fluorescence volumetric imaging. Taken together, these strategies highlight the promise of measurement-efficient volumetric imaging, yet they remain constrained by acquisition speed, optical sectioning, encoding generality, and practical implementation for 4D *in vivo* imaging of whole organs.

Beyond these limitations, high-speed volumetric imaging is further challenged by rolling-shutter distortions inherent to widely used scientific CMOS (sCMOS) cameras, which most existing methods do not explicitly model or correct. Although sCMOS sensors offer an attractive balance of speed, quantum efficiency, and accessibility for dynamic imaging, their line-by-line exposure introduces temporal offsets between adjacent lines. When specimen motion or scanning occurs on timescales comparable to, or faster than the exposure interval, these artifacts manifest as severe rolling-shutter distortions^30^. Such distortions degrade morphology, confound motion estimation, and bias quantitative analysis. Some sCMOS sensors offer a global-shutter mode to eliminate these effects, but doing so typically reduces achievable frame rates, increases readout noise, or requires complex synchronization schemes^31^. Other specialized sensors, including ultra-high-speed cameras^8,32^ and event-camera architectures^33^, have higher effective bandwidth. However, their sensitivity and noise characteristics often pose challenges for photon-starved *in vivo* fluorescence imaging, and their adoption is further restricted by cost and limited availability.

In this context, these challenges motivated us to develop compressive axial-integrated planar scanning (CAPS) microscopy, which combines rapid light-sheet scanning with detection-side optical multiplexing to improve measurement efficiency while preserving optical sectioning and temporal fidelity for high-speed volumetric imaging of fast 4D dynamics. CAPS applies a compressed-sensing strategy to encode multiple axial planes within each camera exposure, enabling reconstruction of volumetric dynamics from substantially fewer measurements than required for fully sampled acquisition. The hardware implementation couples galvo-driven light-sheet scanning on the illumination path with electrically tunable lens (ETL)-based remote focusing and pseudo-random encoding masks generated by a digital micromirror device (DMD) on the detection path. Motivated by compressed sensing^34^, which benefits from incoherent measurements, these masks introduce randomized encoding to promote measurement incoherence for reconstruction. During each exposure, fluorescence from multiple encoded planes is multiplexed into a single frame. Volumes are then reconstructed using a plug-and-play alternating direction method of multipliers (PnP-ADMM) scheme that couples a data-consistency term with a denoising-based implicit prior to suppress artifacts and noise. Additionally, although CAPS is compatible with both global- and rolling-shutter captures, synchronizing the DMD with rolling-shutter readout stamps each sensor line with its corresponding encoding pattern. After demultiplexing the raw frames into reconstructed images, lines can be selectively recombined according to their time stamps, thereby improving temporal alignment and mitigating rolling-shutter distortion.

Our results demonstrate that CAPS acquisition and reconstruction substantially increase the effective use of detector readout bandwidth, yielding more than 7500 effective frames (1920 × 404 pixels) per second (fps), corresponding to a 15-fold improvement in imaging throughput relative to the same system operated without CAPS. For 4D cardiac imaging, CAPS achieved 200 volumes per second (vps) over a 100-µm axial range, or 100 vps over a 190-µm range. Decorrelation-based analysis^35^ indicates lateral and axial resolutions of approximately 2.1 µm and 4.5 µm, respectively, enabling organ-level investigations with cellular-scale resolving power. Imaging of fluorescent beads and quiescent zebrafish hearts further demonstrated improved image fidelity after correcting rolling-shutter distortion, with median structural similarity index (SSIM) increasing from 0.77 (interquartile range, IQR = 0.02) to 0.86 (IQR = 0.02; n = 200). Leveraging the enhanced imaging throughput enabled by widely accessible devices and robust computational reconstruction, we applied the integrated CAPS framework to *in vivo* investigations of cardiac contractile function and intracardiac flow in zebrafish larvae. These capabilities support beat-resolved quantification across scales, spanning organ-level analyses of chamber-volume dynamics and rhythm perturbations to cellular-level tracking of heterogeneous myocardial motion and intracardiac blood-flow patterns under both physiological and pathophysiological conditions.

Collectively, CAPS improves detector-bandwidth utilization, preserves high-speed optical sectioning, and mitigates rolling-shutter distortion. Combined with continuing advances in computational hardware and algorithms, these capabilities position CAPS to expand volumetric imaging and analysis throughput and to support system-level studies of biological dynamics at micrometer resolution and millisecond timescales in both health and disease.

## Results

### Principle and Design of CAPS Microscopy

CAPS hardware system integrates scanned light-sheet excitation, synchronized detection, and mask-encoded readout to achieve high-throughput volumetric imaging with rolling-shutter distortion correction (**Supplementary Note 1, Supplementary Figs. 1–2**). A galvanometer driven by a sinusoidal waveform scans the excitation light sheet along the detection axis. The sinusoidal drive minimizes abrupt accelerations, thereby reducing mechanical resonance and inertia-induced lag^7^. To compensate for the non-uniform axial sampling density and intensity weighting introduced by the sinusoidal scan trajectory, we resample the data onto a uniform axial grid and apply intensity normalization prior to quantitative analyses. An ETL conjugated to the back focal plane of the detection objective shifts the detection focus in synchrony with the light-sheet scan. Operating at 100 Hz with forward and backward sweeps, the system provides high-speed volumetric acquisition at 200 vps (**Fig. 1a**). Precise alignment between the illumination plane and detection focus throughout the scan is achieved using an image-based calibration workflow built upon the Shannon entropy of the normalized discrete cosine transform (DCTS) metric^36^, ensuring optimal optical sectioning (**Supplementary Note 1, Supplementary Fig. 3**). Optical performance after alignment is validated across the operating range using PSF measurements acquired at multiple ETL-galvo synchronization frequencies, demonstrating a lateral resolution of 2–3 µm and an axial resolution of 2–4 µm, prior to compressive acquisition (**Supplementary Figs. 4–5**). Fluorescence emission is encoded by a DMD positioned at the intermediate image plane and subsequently recorded by an sCMOS camera. DMD and camera are pixel-wise registered for precise encoding (**Supplementary Fig. 6**). The sCMOS sensor serves as the master clock, issuing an exposure-start trigger that initiates, for each rolling-shutter exposure, a predefined sequence of binary DMD masks, with the sequence length defined as the compression ratio (CR) (**Fig. 1b, Supplementary Figs. 7–8**).

**Fig. 1.**
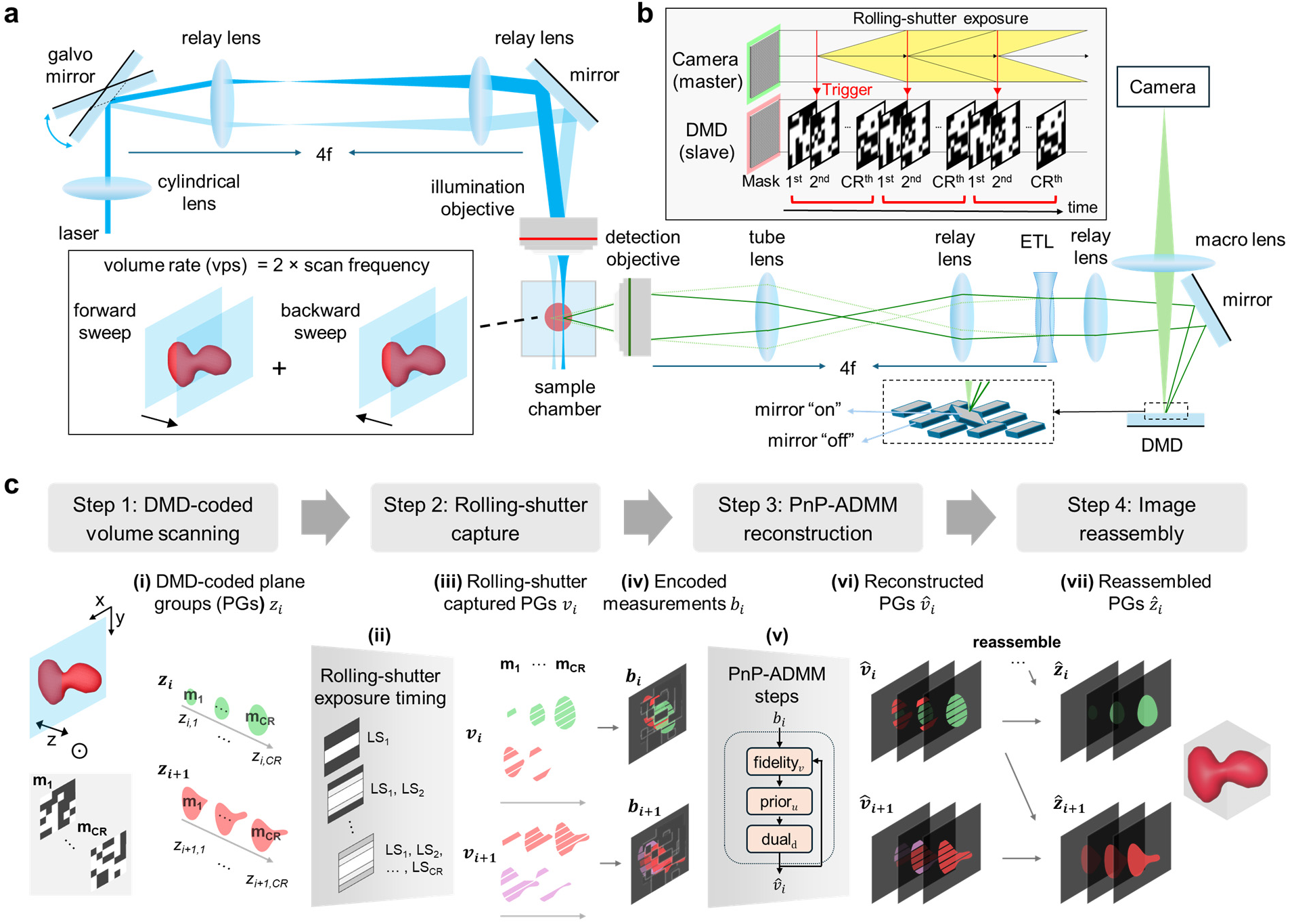
Principle and design of CAPS microscopy. **a,** Optical layout of CAPS. A galvanometer scans the excitation light sheet along the detection axis while an ETL synchronously shifts the detection focal plane; bidirectional sweeps are used to maximize volume rate. Light reflected from the “on” micromirrors is collected by the camera. **b,** Timing synchronization. The sCMOS camera acts as the master clock; each exposure-start trigger initiates a deterministic DMD mask cycle (slave) within the rolling-shutter exposure window. **c,** CAPS processing pipeline that comprises four main steps: DMD-coded volumetric scanning, rolling-shutter capture, PnP-ADMM reconstruction, and image reassembly to form the reconstructed volume (i–vii). **(i)** During continuous axial scanning, the sweep is discretized into plane groups *z_i_* (and *z_i+1_*), each containing CR axial planes *z_i,r_*, and fluorescence within each group is encoded by a sequential DMD mask cycle (m_1_ to m_CR_). **(ii)** Rolling-shutter capture produces staggered line exposure; ordered line segments are denoted LS_1_ to LS_CR_ (earliest to latest exposure timing). **(iii)** The resulting line-dependent mixtures are represented as plane groups *v_i_* (and *v_i+1_*), with line-to-time mapping determined by the calibrated rolling-shutter timing relative to DMD mask timestamps. **(iv)** These multiplexed mixtures are integrated into raw camera frames *b_i_* (and *b_i+1_*). **(v)** For each exposure, PnP-ADMM reconstruction solves *b_i_ = Av_i_ + η* to recover *v̂ _i_* from *b_i_*, including alternating data-fidelity, prior, and dual updates. **(vi)** The reconstructed plane groups *v̂ _i_* and *v̂ _i+1_* retain the line-wise rolling-shutter timing signature. **(vii)** Using calibrated line timing, line segments from *v̂ _i_* and *v̂ _i+1_* are reassembled to restore the plane groups *ẑ_i_* and *ẑ_i+1_*.

In CAPS, fluorescence from multiple axial positions is encoded into each individual camera frame. The relative contribution of each axial position is deterministically set by the line-dependent timing of the sCMOS rolling-shutter exposure together with the calibrated DMD mask sequence (**Supplementary Fig. 9**). Reconstruction therefore combines PnP-ADMM-based recovery of the encoded fluorescence signals with line-timing-guided reassembly to correct rolling-shutter distortion (**Fig. 1c, Supplementary Video 1**). The workflow comprises four stages. First, during continuous axial scanning, the sweep is discretized into plane groups *z_i_*, where *i* indexes the groups acquired over the course of the scan, and each group contains CR axial planes *z_i,r_* (*r = 1,…,CR*). Fluorescence from the planes within each group is encoded by a corresponding set of DMD masks *m_1_* to *m_CR_* (**Fig. 1c(i)**). Second, because the sCMOS sensor exposes lines at staggered times while scanning and coding continue, different line segments within a single exposure sample different subsets of encoded planes. We denote these ordered line segments as *LS_1_* to *LS_CR_*, from earliest to latest exposure timing (**Fig. 1c(ii), Supplementary Fig. 10**). Their combined line-dependent fluorescence mixture forms a rolling-shutter captured plane group *v_i_*. The correspondence between line segments and mask timing is determined from the calibrated DMD timestamps relative to the rolling-shutter exposure (**Fig. 1c(iii)**) and is recorded in the raw camera frame *b_i_* (**Fig. 1c(iv)**). Third, *v_i_* is reconstructed from *b_i_* as *v̂ _i_* by solving an inverse problem using the PnP-ADMM reconstruction framework^37^ (**Supplementary Note 2)**. The algorithm alternates updates of the data-fidelity term, the prior term and the dual variable, thereby decoupling the forward model from the prior for efficient reconstruction (**Fig. 1c(v, vi)**). We present experimental results using the total variation prior^38^, which is shown to be effective. Fourth, because *v̂ _i_* remains in rolling-shutter timing coordinates, we use the calibrated line-timing information to reassign line segments into their correct mask-time bins and combine complementary segments across adjacent plane groups to recover reassembled planes (**Supplementary Note 3, Supplementary Fig. 11**). For example, complementary line segments in *v̂ _i_* and *v̂ _i+_*_1_ are reassembled to restore the plane groups *ẑ_i_* and *ẑ_i+1_* (**Fig. 1c(vii)**). The reassembled plane groups are then integrated into the volume stack. As a result, CAPS increases the effective axial sampling throughput by a factor of CR relative to one-plane-per-exposure acquisition and enables computational correction of rolling-shutter distortion.

### Quantitative Assessment of CAPS Performance

We characterized CAPS imaging strategy using *in silico* 4D beating-phantom models, retrospective data analysis, and integrative hardware-algorithm performance assessment (**Fig. 2a**), quantified by SSIM, peak signal-to-noise ratio (PSNR), normalized root-mean-square error (NRMSE), and decorrelation-based resolution estimation^35^. To assess the reconstruction fidelity of 4D dynamics using the PnP-ADMM framework, we built a synthetic two-chamber beating model that simulates periodic cardiac contraction and performed *in silico* compressed measurements (CR = 15) across 50 volumes spanning a full cycle (**Fig. 2b, Supplementary Video 2**). Chamber motion was driven by antiphase sinusoidal functions to mimic reciprocal contraction and relaxation, with individual cell diameters set to 5–7 µm. Quantitatively, pairwise comparisons between the reconstructed volumes and ground truth (GT) yielded a median PSNR (IQR) of 32.22 dB (0.08 dB), and a median SSIM (IQR) of 0.94 (0.01) (**Fig. 2c**). Median and IQR were used to robustly summarize the time-resolved volumes. The reconstructed 4D chamber volumes closely tracked the GT, with overall NRMSE values of 0.05 and 0.04 for the two chambers, respectively (**Fig. 2d**). These results indicate that the PnP-ADMM reconstruction workflow accurately recapitulates both global and regional spatiotemporal changes.

**Fig. 2.**
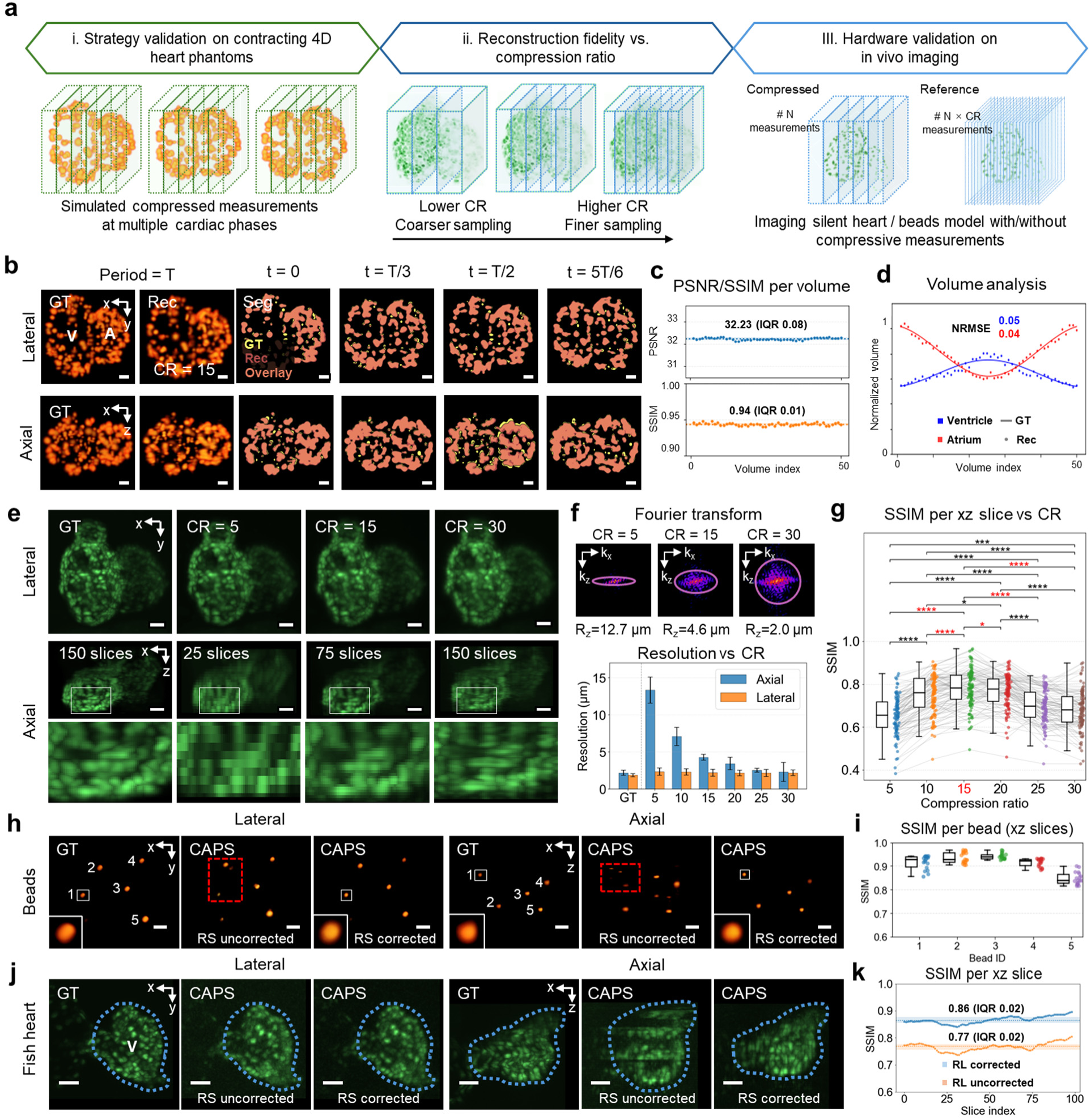
Evaluation of CAPS performance. **a**, Overview of the evaluation workflow combining synthetic modeling, retrospective data analysis, and hardware-algorithm benchmarking. **b**, Comparison of GT and CAPS reconstruction (Rec) at CR = 15 for a synthetic two-chamber 4D beating phantom (V, ventricle; A, atrium). Lateral and axial maximum-intensity projections (MIPs) of the binarized GT volume and CAPS reconstruction are overlaid in distinct colors to visualize their spatial correspondence at representative time points across the beating cycle. Scale bar, 30 µm. **c, d,** Quantitative phantom fidelity across 50 volumes: PSNR/SSIM (**c**) and chamber volume-time tracking (**d**) for the two chambers with NRMSE analysis. **e**, Lateral and axial MIPs of retrospective zebrafish heart reconstructions for the GT and CR = 5, 15, 30, illustrating differences in effective axial sampling density (boxed regions highlight sampling- and fidelity-dependent details). Scale bar, 30 µm. **f,** Top, Fourier transforms of the central xz slice at CR = 5, 15, and 30, with the estimated cutoff frequency indicated by a circle, showing that increasing CR extends the recoverable axial spatial frequency range and supports finer axial detail. Bottom, lateral and axial resolution across CRs, shown as mean ± SD (n = 90). **g**, Reconstruction fidelity versus CR with matched xz slices: SSIM is shown as box plots (median and IQR) with individual slice values overlaid (100 depth-matched slices). Statistical significance across CR groups was assessed using a Friedman test with slice as the blocking factor (𝜒_2_(5) = 300.40, 𝑃 < 0.001, Kendall’s 𝑊 = 0.601), followed by paired Wilcoxon signed-rank tests with Holm correction for post hoc comparisons. A moving block bootstrap sensitivity analysis (block length 3 to 5 slices) yielded the same overall conclusion, supporting robustness to depth-wise slice correlation. Asterisks indicate Holm-corrected pairwise significance (*𝑃_Holm_ < 0.05, **𝑃_Holm_ < 0.01, ***𝑃_Holm_ < 0.001, ****𝑃_Holm_ < 0.0001). Red asterisks denote pairwise comparisons involving the CR of 15. **h,** Bead imaging benchmark: comparison of reference axial-scan volumes (GT) with CAPS reconstructions before and after rolling-shutter correction (boxed regions highlight geometric artifacts). **i,** Bead shape fidelity quantified by SSIM across beads and axial slices. Scale bar, 30 µm. **j,** Quiescent *Tg(myl7:nucGFP)* zebrafish larval heart (3 dpf, *tnnt2* morpholino) acquired with CAPS at 100 vps over a 190-µm axial range: fully sampled motorized axial-scan reference (GT, 1-µm steps) compared with CAPS reconstructions without and with rolling-shutter correction; chamber contours are overlaid for visual comparison (V, ventricle). Scale bar, 30 µm. **k,** Rolling-shutter correction improves per-slice SSIM along the axial direction in the quiescent heart dataset.

To investigate the contribution of CR to spatial resolution and reconstruction fidelity under realistic noise conditions, we used a static 3D volume comprising 150 axial slices extracted from retrospectively synchronized transgenic *Tg(myl7:nucGFP)* zebrafish hearts as GT^11^. We simulated compressed measurements by encoding this volume with predefined (pseudo-random) binary masks at CR values of 5, 10, 15, 20, 25 and 30 (examples for CR = 5, 15, 30 are shown in **Fig. 2e**). Given the constraints of 500 fps acquisition (corresponding to a 404-line readout on a Hamamatsu Flash 4.0 v3 camera) and a target minimal rate of 100 vps for whole-heart imaging, we fixed the acquisition to five compressed measurements per cardiac volume, and reconstructed datasets containing 25, 50, 75, 100, 125 and 150 slices corresponding to the respective CR values. Before generating compressed measurements, the GT volume was axially binned to match the corresponding reconstruction voxel size at each CR. Spatial resolution for each reconstruction was estimated using the effective optical transfer function cutoff frequency derived from decorrelation-based analysis (**Supplementary Note 4, Supplementary Fig. 12**)^35,39^. The Fourier transform of an axial (xz) center slice for CR = 5, 15, and 30, together with the estimated spatial cutoff frequency, illustrated the resolution trends (**Fig. 2f, top**). Increasing CR from 5 to 30 produced a monotonic increase in the axial cutoff frequency, corresponding to a decorrelation-based resolution estimate that improved from ∼14.1 µm to ∼2.3 µm (**Fig. 2f, bottom**). Although this trend is consistent with finer axial discretization at higher CR, the reduced measurement support per slice at higher CR means that this metric should be interpreted as an estimated axial frequency trend rather than a direct claim of validated resolving performance. To assess reconstruction fidelity across CRs while preserving axial structural content, we quantified the similarity between GT and reconstructed axial slices. Because axial sampling density varies with CR, reconstructed volumes were matched to GT dimensions by voxel replication along the detection axis (z-axis), avoiding smoothing artifacts induced by interpolation. Median slice-wise SSIM values, reported as median (IQR) to account for depth-dependent structural heterogeneity, increased from 0.656 (0.119) at CR = 5 to 0.760 (0.133) at CR = 10, peaked at 0.783 (0.112) at CR = 15, and then decreased to 0.776 (0.103) at CR = 20, 0.696 (0.117) at CR = 25, and 0.681 (0.120) at CR = 30. A Friedman test with slice as the blocking factor showed a significant effect of CR on SSIM (𝜒_2_(5) = 300.40, P < 0.001), indicating that reconstruction fidelity varied systematically with CR rather than arising from random slice-wise variation. Post hoc paired comparisons further showed that CR = 15 differed significantly from the other CR conditions (**Fig. 2g**). PSNR showed a similar intermediate optimum (𝜒_2_(5) = 188.93, P < 0.001), with the highest median value at CR = 20 (25.71 dB) and comparable performance at CR = 15 (25.07 dB), indicating a high-performance regime spanning CR from 15 to 20 (**Supplementary Fig. 13**). Together, these results reveal an intermediate CR optimum that reflects the trade-off between improved axial representation at higher CR and the increasing ill-posedness and noise sensitivity imposed by a fixed compressed measurement budget, consistent with expectations from regularized inverse problem theory and compressed sensing under-sampling limits^21,40,41^. We selected CR = 15 for subsequent zebrafish experiments, as it yielded the highest SSIM among the matched axial slices while providing a more robust operating point.

To evaluate reconstruction fidelity after CAPS acquisition and rolling-shutter correction across both controlled and biologically realistic structures, we imaged fluorescent beads (∼10–14 µm in diameter) (**Fig. 2h, i**) and a quiescent heart from a transgenic *Tg(myl7:nucGFP)* zebrafish larva at 3 days post-fertilization (dpf) treated with *tnnt2* morpholino (**Fig. 2j, k**). Both specimens were acquired using CAPS at 100 vps over a 190-µm axial range, followed by fully sampled reference volumes obtained via motorized z-scanning at 1-µm steps. In both cases, CAPS successfully captured and reconstructed the salient features of individual beads and cardiomyocyte nuclei. However, without rolling-shutter correction, systematic distortions were apparent: bead reconstructions exhibited pronounced geometric artifacts (highlighted by red boxes in **Fig. 2h**), and heart images (**Fig. 2j, Supplementary Video 3**) displayed distinct contour discrepancies in the ventricle relative to GT references. Because rolling-shutter correction primarily improves contour and structural agreement with the reference, we focused on SSIM in this comparison. Quantitatively, bead reconstructions showed high slice-wise structural similarity after correction, with bead-wise median SSIM (IQR) values of 0.9266 (0.0428), 0.9263 (0.0409), 0.9399 (0.0171), 0.9211 (0.0281), and 0.8385 (0.0374) across 18 axial slices per bead (n = 5 beads; **Fig. 2i**). For the morpholino-treated heart, median slice-wise SSIM increased from 0.77 (0.02) to 0.86 (0.02; n = 200) following rolling-shutter correction (**Fig. 2k**). Decorrelation-based analysis^35,39^ of the corrected images indicated a lateral and axial resolution of 2.1 µm and 4.5 µm, respectively. These results demonstrate that CAPS preserves reconstruction fidelity while increasing volumetric imaging throughput across specimens of varying complexity, enabled by the integration of mask-encoded acquisition, PnP-ADMM reconstruction, and rolling-shutter distortion correction.

### CAPS Imaging of Beating Zebrafish Heart

Quantitative *in vivo* measurement of 4D cardiac dynamics at high spatiotemporal resolution is essential for linking organ-level function to cellular-level kinematics. By improving spatiotemporal sampling and data-throughput efficiency, CAPS addresses two central requirements for 4D cardiac imaging and supports an integrated zebrafish cardiac imaging and analysis pipeline. CAPS enables high-speed imaging over a 1920 × 404-pixel field of view (∼1037 × 218 µm^2^) at 7500 fps with a CR of 15, corresponding to an effective pixel rate of 5.82 GHz. For *in vivo* cardiac imaging, the entire beating heart of a representative 3 dpf *Tg(myl7:nucGFP)* zebrafish larva could be captured within a reduced lateral region of interest of ∼313 × 218 µm^2^ and an axial scanning range of 190 µm at 100 vps without rolling-shutter correction (**Fig. 3a**). For this reduced region-of-interest setting, the effective pixel throughput was ∼1.76 GHz, compared with ∼0.12 GHz for native rolling-shutter acquisition without CAPS, and far exceeded the ∼0.06 GHz theoretical throughput achievable with conventional global-exposure synchronization^31^. CAPS also substantially improved data-throughput efficiency, requiring only ∼0.23 GB/s of physical readout to reconstruct a volumetric stream equivalent to ∼3.51 GB/s of uncompressed data (**Fig. 3b**).

**Fig. 3.**
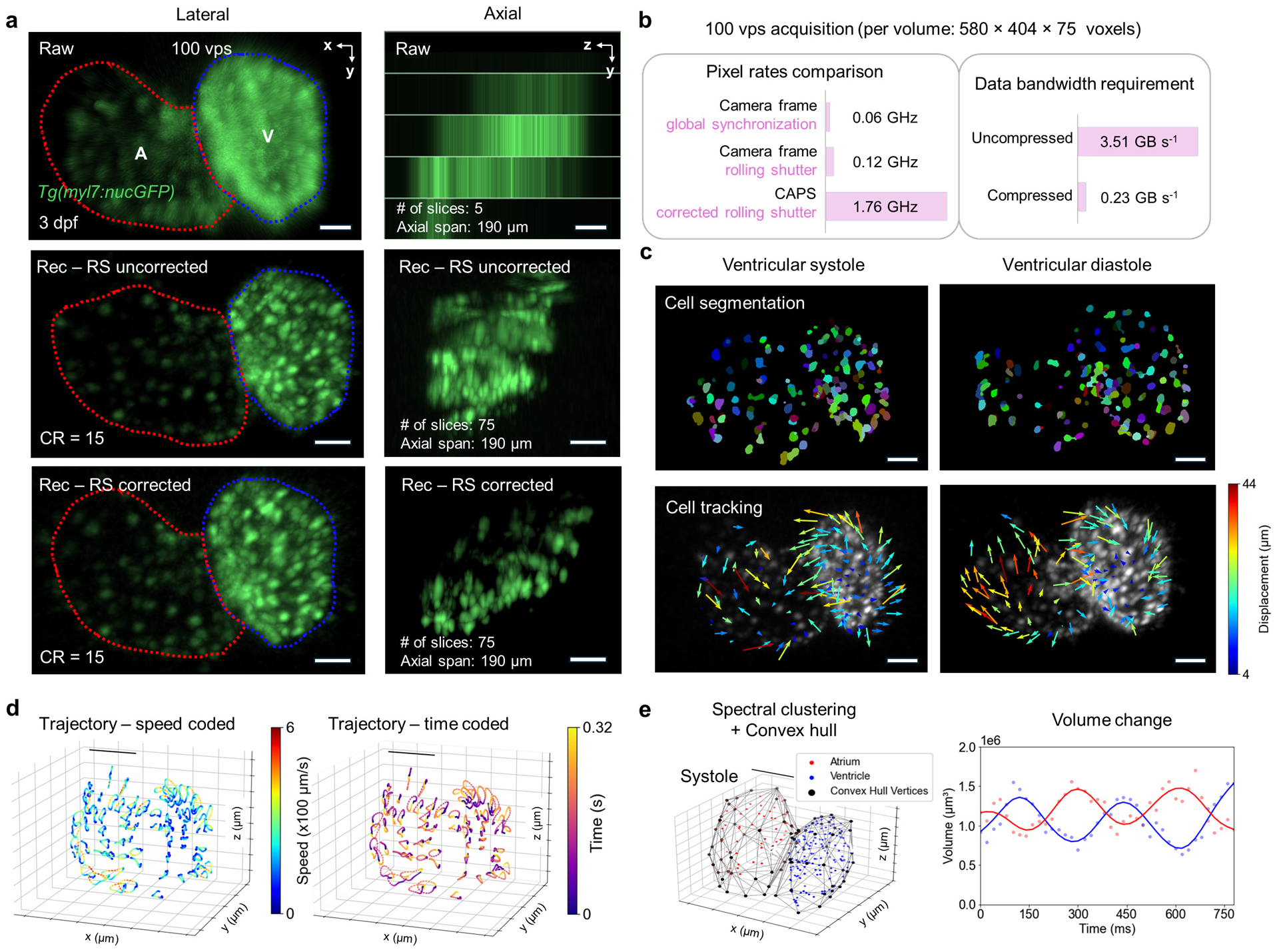
High-speed 4D CAPS quantification of zebrafish heart dynamics. **a**, Representative lateral and axial MIPs of the CAPS raw measurements and reconstructed volumes of a 3 dpf *Tg(myl7:nucGFP)* heart acquired at 100 vps (CR = 15), shown before and after rolling-shutter realignment (V, ventricle; A, atrium). Raw acquisition contains 5 frames over a 190-µm axial range, while reconstruction yields 75 planes over the same range. **b,** System throughput and data efficiency at 100 vps (per volume: 580 × 404 × 75 voxels). Effective pixel rate comparison for global-exposure-synchronized, native rolling-shutter, and CAPS rolling-shutter-corrected output (left), and equivalent uncompressed bandwidth (3.51 GB/s) and compressed physical readout (0.23 GB/s) (right). **c,** Cardiomyocyte nuclei segmentation (top) and tracking (bottom) at ventricular systole and diastole, with motion vectors color-coded by displacement from systole to diastole (left) and from diastole to systole (right). **d,** Single-cell 3D trajectories visualized as speed-coded and time-coded maps across a cardiac cycle. **e,** Chamber assignment by spectral clustering and convex hull-based volume reconstruction from nuclear centroids (left), and the corresponding time-resolved atrial and ventricular volume changes (right).

Using CAPS, we established a framework for imaging and quantifying 4D zebrafish cardiac dynamics without the need for retrospective cardiac synchronization. To quantify cellular-level kinematics, we implemented a computational pipeline utilizing BiaPy^42^ for nuclear segmentation and Ultrack^43^ for multi-cycle cell tracking. This enabled continuous tracking of all resolved cardiomyocytes across successive cardiac cycles (**Fig. 3c** and **Supplementary Video 4**). Speed-coded trajectories revealed clear, cyclic bursts of nuclear motion with peak speeds of ∼320–760 µm/s and mean speeds of ∼140–300 µm/s. Complementary time-coded trajectories preserved instantaneous spatial organization and highlighted coordinated motion patterns throughout the cardiac cycle (**Fig. 3d**). To assess chamber-level dynamics, we classified atrial and ventricular nuclei using spectral clustering^44^, and estimated time-varying chamber volumes via a convex hull-based reconstruction^45^ (**Fig. 3e** and **Methods**). The resulting volumetric traces resolved the expected anti-phase dynamics, including peak ventricular filling immediately following atrial systole, and revealed beat-to-beat variability in both amplitude and timing. In the representative heart, ventricular volume decreased from 1.35 × 10^6^ µm^3^ (end-diastole) to 0.78 × 10^6^ µm^3^ (end-systole), corresponding to a stroke volume of 0.57 × 10^6^ µm^3^ (42.2% ejection fraction). Atrial volume decreased from 1.46 × 10^6^ µm^3^ to 1.02 × 10^6^ µm^3^, yielding a stroke volume of 0.44 × 10^6^ µm^3^ (30.1% volume change).

### Quantitative Imaging of Cardiac Contractile Dysfunction

Cardiac contractile dysfunction is often paroxysmal and asymptomatic until the onset of severe complications, posing significant challenges for retrospective^1,10,11^ or prospective^12^ gating imaging approaches. To demonstrate the capability of CAPS to resolve such transient abnormalities, we performed proof-of-concept imaging in 3 dpf larvae that were either untreated or treated with 10 µM amiodarone for 24 h, an antiarrhythmic known to reduce heart rate and induce atrioventricular (AV) block in zebrafish^46,47^. CAPS resolved the contraction and relaxation phases of both chambers with cellular resolution (**Fig. 4a**). To quantify drug-induced changes in chamber contractility, we extracted atrial and ventricular volume trajectories across consecutive cycles. The amiodarone-treated heart exhibited an alternating pattern in which atrial contractions were not consistently followed by full ventricular volume changes, in contrast to the regular beat-to-beat coupling pattern observed in untreated hearts (**Fig. 4b, Supplementary Video 5**). Fourier amplitude spectra of chamber-volume traces showed that untreated hearts were dominated by a single narrowband peak in both ventricle and atrium, indicating near-periodic dynamics with limited spectral dispersion^48^. Following amiodarone treatment, the spectral amplitude redistributed across multiple frequency components: the ventricle exhibited peaks at 2.68 Hz and 5.36 Hz of comparable amplitudes, while the atrium showed a dominant component at 3.12 Hz with an additional peak at 5.39 Hz. This departure from a single-timescale rhythm, together with the approximate 2× relationship between dominant components, is consistent with a 2:1 AV block^46^ (**Fig. 4c**). This frequency-domain signature provided a quantitative indicator of beat-resolved, multi-scale chamber dynamics captured by CAPS, offering a scalable readout for drug-induced abnormalities in cardiac rhythm and supporting rapid screening of dysfunctional phenotypes^49^.

**Fig. 4.**
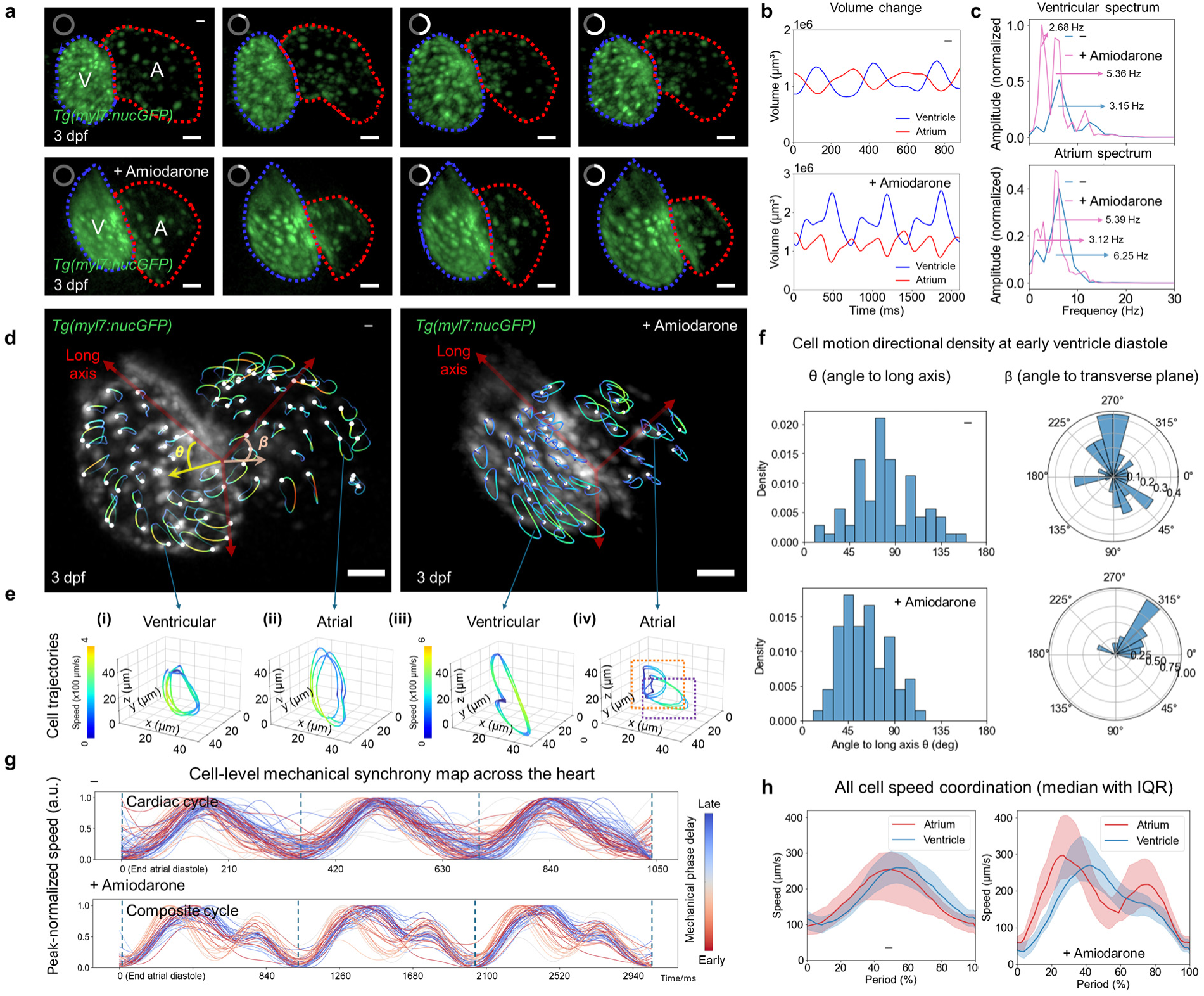
CAPS detects cellular desynchrony and chamber-scale dysfunction in drug-treated hearts. **a,** Beat-resolved whole-heart reconstructions of *Tg(myl7:nucGFP)* larvae at 3 dpf imaged at 100 vps, comparing hearts without treatment and after 24 h exposure to 10 µM amiodarone (V, ventricle; A, atrium). The circumferential markers at the top left of each panel indicate representative phases of the cardiac cycle. For the heart without treatment, markers denote end-systole, ventricular filling, end-diastole, and ventricular ejection. For the amiodarone-treated heart, markers denote key phases of the altered atrioventricular filling pattern, in which atrial contractions were not consistently followed by full ventricular volume change, including end-systole, ventricular filling, atrial filling, and end-diastole. **b,** Atrial and ventricular volume changes across consecutive cycles, showing regular alternation without treatment and disrupted atrial-to-ventricular mechanical coupling under amiodarone treatment. **c,** Fourier amplitude spectra of chamber-volume traces, showing a transition from a dominant narrowband peak in untreated hearts to redistributed spectral amplitude across multiple frequency components after drug treatment. **d,** Whole-heart single-nucleus tracking with instantaneous speed derived from 3D trajectories; ventricular long-axis coordinate system used to define *θ* and *β*. **e**, Representative ventricular and atrial trajectories illustrating repeatable loops without treatment and alternating loops under amiodarone (purple and orange boxes indicate the first and second atrial loops, respectively). **f**, Directional distributions of instantaneous cell displacement at the onset of ventricular diastole, plotted as *θ* histograms and *β* polar maps, revealing a broad, diffuse pattern without treatment and a more concentrated, lobe-dominated pattern under amiodarone. **g**, Phase-colored, peak-normalized single-cell speed traces showing coordinated timing across three consecutive cycles without treatment and broadened phase dispersion under amiodarone; a two-beat composite cycle is used to visualize alternating strong and weak atrial contraction. **h,** Chamber-scale speed distribution summaries of phase-aligned cardiomyocyte speed using the median and IQR, quantifying timing shifts and increased atrial variability under amiodarone relative to ventricular dynamics. Scale bars: 30 µm.

To elucidate motion of the resolved cells across the entire dysfunctional heart, we defined the ventricular long axis (base-to-apex) as a reference for downstream analysis and constructed two orthogonal axes spanning the corresponding transverse plane^50^. The polar angle *θ* denotes the orientation relative to the long axis, and the azimuthal angle *β* specifies position within the transverse plane referenced to the atrium-facing direction of the ventricle (**Fig. 4d**). In the untreated heart, nuclei followed smooth, repeatable 3D trajectories with consistent peak speeds over three consecutive cycles (representative cells in **Fig. 4e(i, ii)**), with ventricular peak velocity of ∼228 µm/s and atrial peak velocity of ∼241 µm/s. In contrast, the amiodarone-treated heart exhibited kinematic irregularity. The representative ventricular cell completed only a single closed trajectory loop (peak velocity ∼304 µm/s) over a time interval corresponding to two nominal cycles, whereas the atrial cell executed two distinct loops (purple vs orange) with peak velocities of ∼261 µm/s and ∼164 µm/s, respectively (**Fig. 4e(iii, iv)**). In addition to velocity profiles, we investigated the geometry of collective 3D motion across the chamber. For each nucleus, we computed the instantaneous displacement direction and expressed it in spherical coordinates using the angles *θ* and *β* (**Fig. 4f**). The resulting angular histograms reflected directional bias and alignment, following conventions for analyzing biological movement vectors and circular data^51,52^. At the onset of ventricular diastole (pooled across three cycles), the untreated heart exhibited a broadly distributed set of displacement directions, with a wide θ distribution and a diffuse, multilobed *β* pattern (**Fig. 4f, top**), indicating weak directional concentration^52,53^. This dispersion aligns with the mechanical complexity of early diastolic relaxation, during which myocardial restoring forces and elastic recoil drive rapid, spatially heterogeneous deformation^54^. In contrast, the amiodarone-treated ventricle showed a more concentrated directional distribution, with *θ* values clustering toward oblique orientations and the *β* polar plot collapsing into a dominant lobe (**Fig. 4f, bottom**), suggesting reduced spatial heterogeneity and altered organization of diastolic motion^52,53^.

Beyond directionality, CAPS also revealed the temporal sequence of cardiomyocyte activation by identifying when individual cells reached their peak contractile activity within each cycle. This enabled direct comparison of coordinated contractile wave propagation and chamber-level dynamics under different physiological and pharmacological conditions. From each tracked trajectory, we computed instantaneous speed and peak-normalized the speed profile to emphasize the timing of motion within the cardiac cycle (**Fig. 4g**). In the amiodarone-treated heart, we show three composite cycles, each spanning one pair of alternating strong and weak, incomplete atrial or ventricular contractions. Each speed trace was color-coded by its mechanical phase delay within the normalized cardiac period (phase 0 defined at ventricular end-systole), with red indicating earlier-phase motion and blue indicating later-phase motion. In the untreated heart, beats were highly reproducible and largely phase-locked across three consecutive cycles. Subtle delays in the onset of contractile motion were readily captured by CAPS and were evident among neighboring cells. In contrast, the amiodarone-treated heart showed alternating atrial-to-ventricular coordination consistent with the chamber-level phenotype, revealing the characteristic alternation between strong and weak mechanical events. Notably, within each composite period, the traces segregated along a timing continuum, with trajectories nearer the red end tending to display two prominent speed peaks and those nearer the blue end tending to exhibit a single dominant peak. This separation reflects region-specific responses of atrial and ventricular cells under fixed-ratio rhythm perturbations^46,47^.

To quantify shifts in temporal coordination and heterogeneity among cells, we overlaid three consecutive (or composite) cycles and summarized the cardiomyocyte speed distributions in each chamber over a phase-normalized cardiac period using the median and IQR (**Fig. 4h**). The median trace captures the typical phase-dependent speed profile, whereas the IQR band reports the spread of speeds across cells at each phase. In the untreated heart, both the atrium and ventricle exhibited a single speed peak near mid-cycle (∼50% phase), with median peak speeds of ∼255–260 µm/s. The IQR bands remained relatively narrow throughout the cycle, consistent with comparatively uniform mechanical output across cells. In the amiodarone-treated heart, the atrial population displayed two distinct peaks: an early peak near ∼27% phase (median ∼297 µm/s; IQR ∼173 µm/s) and a later peak (median ∼217 µm/s; IQR ∼129 µm/s). The ventricle retained a single peak near ∼42% phase with a cycle-averaged IQR of ∼67 µm/s. The increased variability in atrial mechanical output, coupled with comparatively preserved ventricular variability, suggests that amiodarone disrupts atrial-to-ventricular mechanical-coupling coordination^55^.

### Quantitative Imaging of Intracardiac Hemodynamics

To investigate hemodynamics arising from rapid, spatially distributed, and often cycle-to-cycle variable flow patterns, we imaged and tracked erythroid cells in *Tg(gata1a:dsRed)* zebrafish larvae from 3 to 4 dpf. This enabled beat-resolved 4D analysis of intracardiac flow, linking transport organization to hemodynamic function. CAPS captured erythroid cells at 200 vps across an axial range of ∼100 µm in 4 dpf larvae (left: raw capture; right: reconstructed volume, **Fig. 5a**), where the more developed heart supports faster and more directional intracardiac transport. We tracked trajectories of fully resolved erythroid cells over consecutive cycles (**Fig. 5b, left**) and quantified their instantaneous speeds (**Fig. 5b, right**). The resulting trajectories revealed diverse intracardiac transport behaviors, including forward transit through the AV canal, brief retrograde excursions consistent with regurgitant flow during ventricular contraction, and prolonged ventricular residence prior to ejection through the bulbus arteriosus **(Supplementary Video 6).** These flow features may contribute to endocardial mechanotransduction and valve morphogenesis in zebrafish, providing an entry point for dissecting the interplay between cardiac pumping dynamics and intracardiac transport^56–58^.

**Fig. 5.**
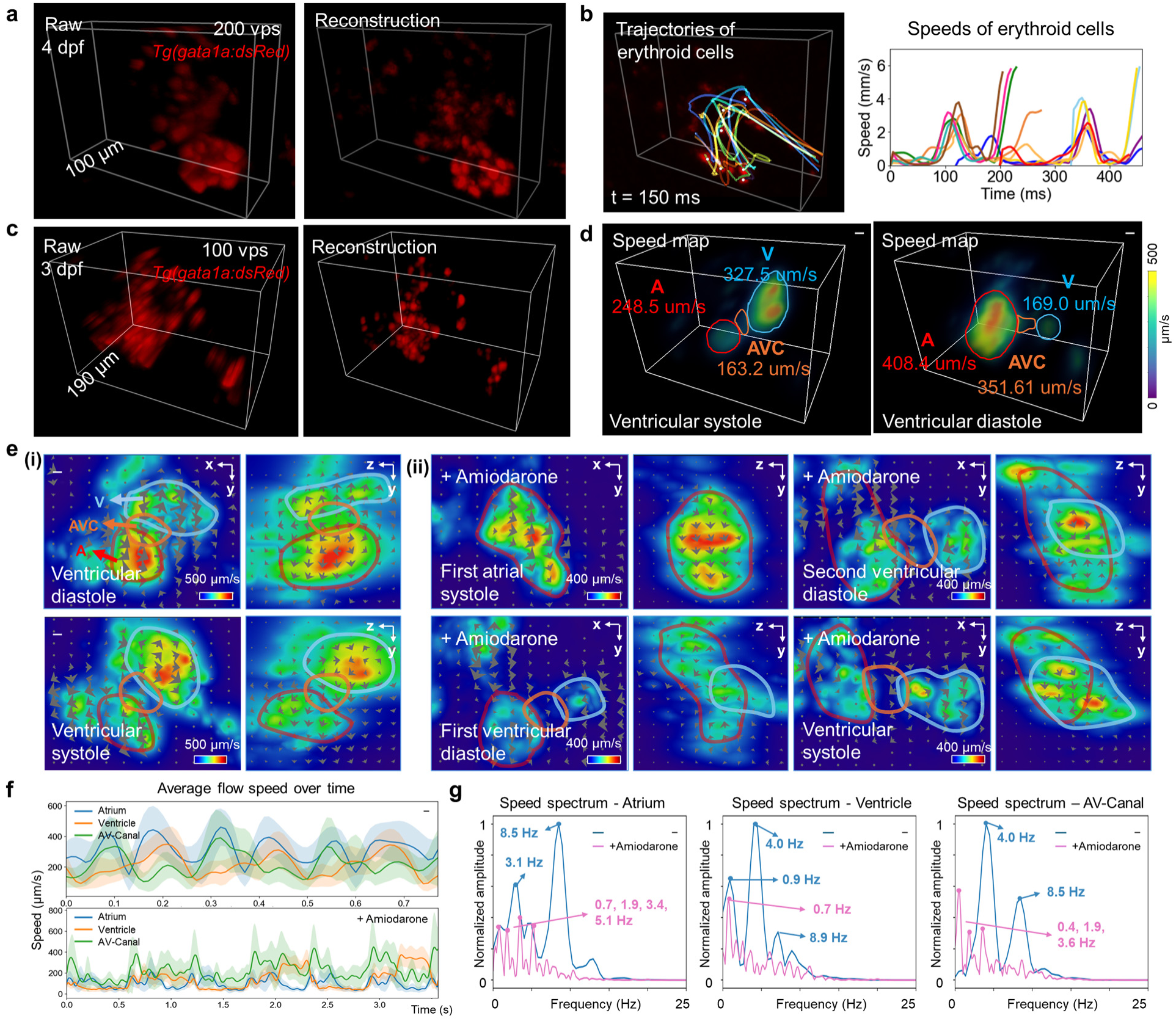
Beat-resolved 4D imaging of intracardiac hemodynamics with CAPS. **a,** CAPS acquisition at 200 vps in a *Tg(gata1a:dsRed)* larval heart, showing the raw measurement (left) and the reconstructed volume (right) at ventricular end-diastole, which resolves individual erythroid cells over an axial range of ∼100 µm. **b,** Example 3D erythroid cell trajectories overlaid on the reconstructed volume (left) with the corresponding speed traces (right), illustrating beat-resolved single-cell transport. **c,** Whole-heart CAPS acquisition at 100 vps over an axial range of ∼190 µm, showing the raw capture (left) and reconstructed volume (right) spanning atrium, AV canal, and ventricle. **d,** Flow velocity magnitude maps at ventricular diastole and systole, with regions of interest for atrium (A, red), AV canal (AVC, orange), and ventricle (V, blue), highlighting compartment-specific inflow and outflow organization. **e,** Phase sequence of CAPS vector-field maps comparing hearts with and without amiodarone, where the treated heart exhibits two atrial inflow activations followed by a delayed ventricular ejection over a composite cycle. **f,** Regional mean speed traces over three cycles in the atrium, AV canal, and ventricle, showing reproducible phase-locked peaks without treatment and altered atrial-to-ventricular coupling with treatment. **g,** Fourier amplitude spectra of the regional mean speed traces for atrium, ventricle, and AV canal. Curves show normalized spectral amplitude computed from the time-domain speed traces in **f**, with peak frequencies annotated. Without treatment, spectra are narrowband with dominant peaks near 3.1 Hz in the atrium and 4.0 Hz in the ventricle and AV canal, plus higher-frequency components near 8.5–8.9 Hz. With amiodarone, power redistributes across multiple components, including atrial peaks near 0.7, 1.9, 3.4, and 5.1 Hz, ventricular power dominated near 0.7 Hz, and AV-canal components near 0.4, 1.9, and 3.6 Hz.

We applied CAPS to volumetric imaging for intracardiac hemodynamic analysis across the atrium, ventricle, and AV canal in 3 dpf larvae at 100 vps over an axial range of ∼190 µm (left: raw capture; right: reconstructed volume, **Fig. 5c**). We estimated 4D blood flow velocity fields using an established flow-field registration workflow that optimizes a displacement grid to align consecutive volumes^43^, and depicted the resulting velocity magnitudes at ventricular end-systole and end-diastole to illustrate ejection and filling processes across chambers (**Fig. 5d** and **Supplementary Video 7**). During ventricular ejection, blood velocity magnitudes were estimated at ∼248.5 µm/s in the atrium, ∼163.2 µm/s in the AV canal, and ∼327.5 µm/s in the ventricle. In contrast, during ventricular filling, velocity magnitudes increased to ∼408.4 µm/s in the atrium and ∼351.6 µm/s in the AV canal, while ventricular velocity magnitudes decreased to ∼169.0 µm/s. This phase-dependent redistribution of flow, characterized by elevated atrial and canal transport during filling and higher ventricular speeds during ejection, aligns with prior findings on dynamic intracardiac flow patterns and canal-specific shear cues^56–58^.

In *Tg(gata1a:dsRed)* zebrafish larvae treated with 10 µM amiodarone at 2 dpf and imaged at 3 dpf, CAPS captured perturbed hemodynamics and rhythm abnormalities. Velocity-field analysis revealed flow direction and magnitude across the full cardiac volume, enabling discrimination of antegrade versus retrograde transport and visualization of how circulation shifts between chambers and through the AV canal over time (**Fig. 5e** and **Supplementary Video 8**). These data are presented as a demonstration of CAPS capability under a perturbed condition, rather than as a population-level assessment of biological variability. In the untreated heart, flow fields followed a coordinated sequence in which atrial inflow accelerated during ventricular filling and converged toward the AV canal as a compact antegrade jet, followed by rising ventricular inflow and a transition to outflow-directed high-velocity ejection (**Fig. 5e(i)**). Correspondingly, regional mean-speed traces were highly reproducible and phase-locked, with atrial peaks (red) preceding AV canal (orange) and ventricular peaks (blue), consistent with coordinated chamber coupling that produces stereotyped transvalvular transport and well-defined canal shear dynamics (**Fig. 5e,f**)^64,66^. Fourier amplitude spectra of the regional mean-speed traces revealed a largely narrowband structure, dominated by one or two principal oscillatory components with clear higher-frequency content (**Fig. 5g**), reflecting strongly periodic flow. In contrast, amiodarone treatment produced a multiphase flow field with disturbed atrial-to-ventricular coordination over a composite cycle (**Fig. 5e(ii)**). Accordingly, regional mean-speed traces exhibited two atrial peaks per composite cycle, whereas AV canal and ventricular traces showed a dominant response associated with the larger ejection and reduced modulation during the secondary atrial event (**Fig. 5f**). This pattern indicated weakened and variable atrial-to-ventricular flow coupling, expected to alter the timing and directionality of canal forces that pattern the developing valve^56^. Fourier analysis showed a distribution of spectral amplitude across multiple components and region-specific differences, reflecting multi-timescale variability and the 2:1 AV block observed in the time-domain flow fields (**Fig. 5g**).

## Discussion

We presented CAPS as a framework that alleviates the detector-bandwidth constraint that traditionally limits high-speed 4D fluorescence microscopy. CAPS optically multiplexes multiple axial planes into each sCMOS frame and computationally demultiplexes them through model-based reconstruction, thereby shifting the performance burden from bandwidth-limited hardware to scalable computation. Compared with previous optical encoding strategies, CAPS combines several advantages that are rarely achieved simultaneously in fluorescence volumetric imaging, including improved measurement efficiency, preserved optical sectioning, and high spatiotemporal resolution, all of which are suitable for quantitative analysis. In parallel, CAPS leverages the deterministic timing relationship between the sCMOS readout and the triggered DMD mask sequence, providing a practical strategy to reduce temporal distortion inherent to rolling-shutter acquisition and improve quantitative reliability for fast 4D dynamics.

Integrated with PnP-ADMM reconstruction, CAPS enables beat-resolved 4D investigation of cardiac contractility and hemodynamics, with particular emphasis on perturbed, irregular heartbeats, at cellular resolution in zebrafish larvae, while maintaining temporal fidelity and substantially reducing data management demands through compressive acquisition. CAPS is also expected to reduce cumulative light exposure for *in vivo* imaging, and thus the risk of photodamage or phototoxicity, relative to retrospective gating approaches^1,10,11^. By combining single-cell tracking, flow field analysis, and spectral analysis, CAPS provides a scalable framework for probing how changes in organ-scale dynamics (e.g., cardiac contractile function, heart rate variability, rhythm disturbances, intracardiac hemodynamics) influence cellular-level kinematics (e.g., motion velocity and direction, displacement, trajectory), and vice versa. Following amiodarone treatment, this framework revealed impaired atrial-to-ventricular mechanical coupling, increased cell kinematic variability, altered motion directionality, broader dispersion of peak activity timing, and perturbed flow coordination across chambers. These readouts link global dysfunction and disturbed circulation patterns to local cellular-scale transport and establish a foundation for future automated metrics, including AV coupling indices^47^, regurgitant transport fractions^59^, and oscillatory shear surrogates^60^, relevant to drug and genetic screens targeting mechano- or flow-sensitive developmental pathways.

As with other compressive, model-based reconstruction approaches, the forward model and the choice of denoising prior matched to the sample’s statistics are fundamental determinants of CAPS performance. In practice, mismatches in the forward model, including imperfect mask calibration, residual rolling-shutter timing error, and nonstationary intensity, or mismatches in the prior can be amplified under higher compression, manifesting as structured artifacts. To improve robustness and modularity, we adopted the PnP-ADMM solver that decouples the measurement model from the regularization mechanism^37^. This design allows the prior to be swapped or tuned without redefining the optimization objective, while retaining convergence guarantees under standard assumptions. In this study, we selected a total variation denoiser as it effectively suppresses noise and small intensity fluctuations while preserving sharp cellular boundaries in cardiac datasets^61^. It also requires simpler tuning and substantially lower computational cost than nonlocal patch-based volumetric denoisers such as BM4D^62^. Building on this workflow, we validated system fidelity and resolution limits across a range of CRs. Looking forward, the reconstruction fidelity could be further strengthened by incorporating other priors, such as deep-denoiser priors^63^, transformers^64^, and microscopy-tailored training strategies^65^. Beyond deterministic reconstruction, generative priors^66^ offer a complementary direction for uncertainty-aware recovery from highly compressed measurements. Future implementations could also incorporate joint spatiotemporal reconstruction^67^ to improve temporal consistency across reconstructed volumes, as computational resources become more cost-effective.

CAPS hardware was designed for both rapid scanning and physics-based forward modeling. For volumetric scanning, a synchronized ETL-galvo pair performs remote refocusing and light-sheet sweeping to traverse depth at 100 Hz. To reduce calibration mismatch, the ETL-galvo synchronization and validation pipeline calibrates the relative phase, offset, and amplitude of the sinusoidal drive waveforms by optimizing an image-quality objective based on DCTS^36^ over a full scan period (**Supplementary Note 1**). Building on this foundation, future implementations can further improve robustness through closed-loop acquisition and real-time control^36^, periodic model verification using fiducials^68^, and drift-aware self-calibration^69^. Axial range and scan fidelity can also be enhanced through alternative remote-focusing architectures to support faster axial scanning^70^, improve light efficiency^71^, and reduce scan-dependent aberrations^72^. To ensure accurate forward modeling, the DMD applies predefined binary masks for each sCMOS frame readout, and per-pixel calibration between the DMD and sCMOS sensor ensures the exact spatial correspondence required for a physics-based forward model that enables compressive demultiplexing. A practical limitation, however, is photon efficiency. In single-arm detection, binary DMD masking redirects light into complementary reflection states, such that collecting only one output effectively discards approximately 50% of the available photons. Because fluorescence imaging is constrained by limited photon budgets and shot noise, this loss becomes consequential for dim structures and at high CRs, where a fixed detected photon budget must support demultiplexing of more encoded images. A dual-port detection scheme that captures the complementary DMD output could improve detection efficiency^73^. Together, these hardware and computational advances provide a foundation for continued improvements in CAPS performance and reconstruction fidelity as digital devices and algorithms evolve.

Complementing the compressive reconstruction and calibration strategies discussed above, CAPS converts sCMOS rolling-shutter readout into a controllable timing dimension that can be modeled and corrected. Synchronization of the DMD with rolling-shutter readout allows line-dependent exposure timing to be stamped by the encoding masks. After the raw frames are demultiplexed into reconstructed images, corresponding lines can be selectively recombined according to these time stamps for improving temporal alignment and suppressing rolling-shutter distortion. At 500 fps with a 7.5 kHz mask refresh rate, dividing each exposure into 15 temporal intervals enables a substantial reduction in rolling-shutter artifacts.

However, it remains an approximation to global shutter acquisition rather than a complete recovery. Specifically, the current model treats lines associated with the same mask within a reconstructed image as having a common effective acquisition time, whereas in practice these lines still experience residual rolling-shutter offsets. Future gains may come from embedding line-dependent timing directly into the PnP-ADMM forward model and pairing it with stronger spatiotemporal priors to more accurately model and correct rolling-shutter effects. In parallel, faster modulation and encoding schedules could further refine temporal alignment, with improved photon efficiency helping preserve signal quality as temporal subdivision increases.

Overall, CAPS offers a practical route to high-speed volumetric fluorescence microscopy by shifting a major bottleneck from detector bandwidth to computational encoding and decoding power, while preserving optical sectioning and mitigating rolling-shutter-induced image distortion. In this work, our strategy allows for quantitative whole-heart measurements spanning chamber-scale dynamics and intracardiac blood flow at cellular resolution, all while keeping the physical readout and data burden manageable. Collectively, the CAPS framework, coupled with rapid advances in computational hardware and algorithms, holds significant promise for expanding volumetric imaging and reconstruction throughput, thereby facilitating system-level investigations of instantaneous biological dynamics at cellular resolution in both health and disease.

## Methods

### CAPS hardware

Two lasers at 473 nm and 532 nm (R471003GX and R531003GX, Laserglow) are expanded 5× and passed through a cylindrical lens to form a static light sheet. The beam is reflected from a kHz galvo mirror (GVS211, Thorlabs) that scans ±10° about its pivot. The galvo, scan lens (*f* = 75 mm, CLS-SL, Thorlabs) and tube lens (*f* = 200 mm, ITL200, Thorlabs) are arranged in a 4f telecentric relay, delivering a sinusoidally swept sheet through an illumination objective (4× 0.13 NA, Nikon). The scan amplitude sets the axial excursion (up to 190 µm) while preserving beam waist and Rayleigh length across the field. Fluorescence is collected with a water-immersion objective (20× 0.5 NA, Olympus) and relayed through the tube lens (*f* = 180 mm, TTL180-A, Thorlabs) into a pair of relay lenses (*f_r_* = 100 mm). At the pupil-conjugate plane of the first relay lens sits a gravity-compensated ETL (EL-16-40-GTC-VIS-5D-C, Optotune), whose sealed fluid chamber allows arbitrary orientation without gravitational sag. The tunable lens spans –2 to +3 diopters (*f_ETL_* ≈ –500 mm to +333 mm), shifting the detection image plane by

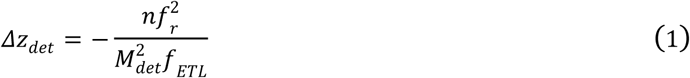

where *n* is the immersion-medium refractive index and *M_det_* is the objective magnification.

Downstream of the ETL, the second relay lens forms the image onto a DMD (DLP9500, Texas Instruments). During each camera exposure, the DMD cycles through multiple CR binary masks, optically encoding sequential axial planes with unique 2D patterns. The relay lenses then project the masked signal onto the sensor. The encoded fluorescence is imaged by a macro lens (AF Micro-NIKKOR 60 mm f/2.8D, Nikon) onto the sCMOS camera (ORCA-Flash 4.0 V3, Hamamatsu) operated with a 1920 × 404-pixel sub-array at 500 fps. The macro lens is set to provide 0.6× lateral magnification, mapping each 10.8 µm DMD micromirror to a 6.5 µm camera pixel. With a CR of 15, the system achieves an effective acquisition rate of 7500 masked frames per second, which corresponds to 100 vps (190-µm range) or 200 vps (100-µm range), respectively. All main system components are listed in **Supplementary Table 1**.

### ETL and galvo synchronization

To maintain axial coincidence between the illumination plane and the detection focal plane during high-speed volumetric scanning, the ETL and galvo are driven by sinusoidal command waveforms, which reduce tracking errors caused by abrupt waveform transitions at 50–100 Hz^7^. For each actuator *j*∈{ETL, Galvo}, the command parameters are defined as *p_j_=(A_j_, Φ_j_, O_j_)*, where *A_j_* is the scan amplitude, *Φ_j_* is the phase, and *O_j_* is the offset. Under this notation, the galvo command waveform is written as

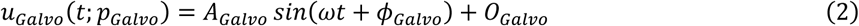

with the ETL defined analogously. The synchronization objective is to identify the galvo command parameters that minimize axial mismatch between the realized illumination plane and detection focal plane while the ETL command is held fixed.

The galvo phase *Φ_Galvo_* is optimized first by sweeping candidate values while keeping *A_Galvo_* and *O_Galvo_* fixed. For each setting, a time series of images {*I_t_*} is acquired over one or more scan cycles, and the calibration cost is defined as

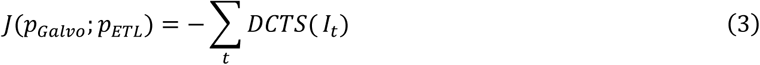

where DCTS is an image-sharpness metric that increases with focus quality and therefore decreases with illumination-to-detection misalignment (**Supplementary Note 1**)^36^. Minimizing *J*(·) is thus equivalent to maximizing cumulative image sharpness over the scan. After identifying the optimal *Φ_Galvo_*, the same procedure is applied sequentially to optimize *O_Galvo_* and then *A_Galvo_*. This phase–offset–amplitude order follows the analytical result that phase mismatch is the most uniquely identifiable contributor to synchronization error, enabling more stable optimization of the remaining parameters.

### PnP-ADMM reconstruction and parallelization

Compressed CAPS frames are reconstructed into volumes using an adapted PnP-ADMM^37^ solver. For clarity, we present here a simplified forward model: *b = Av + η* (the full formulation in **Supplementary Note 2**), where *b* is the 2D camera frame, *v* is the unknown encoded sub-volume, *A* is the known sensing operator that combines axial sweeping with the DMD mask sequence, and *η* represents photon-shot and readout noise. Reconstruction is formulated as

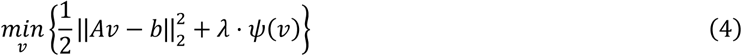

with *ψ(*⋅) a generic image prior. To decouple the data-fidelity term and the regularization term, we introduce an auxiliary variable *u* and rewrite the problem as

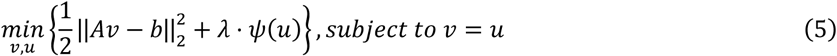

The corresponding augmented Lagrangian is

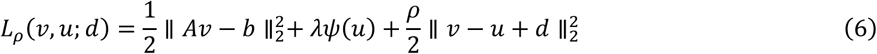

where *ρ* is the ADMM penalty parameter and *d* is the dual variable. The problem is then solved using the following PnP-ADMM updates:

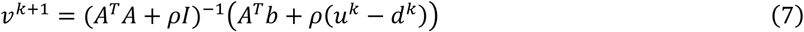

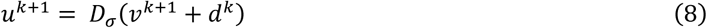

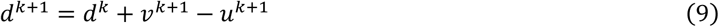

where *D_σ_*(⋅) is an external denoiser, implemented here as total variation denoising, with an estimated noise level σ = √(λ / ρ).

To improve computational efficiency, the reconstruction takes advantage of the block structure inherent to the coded acquisition. Before rolling-shutter correction, each compressed frame can be processed independently in the data-consistency update within each iteration. Per-iteration cost is further reduced by using a closed-form update that leverages the pixel-wise action of the DMD masks, replacing large coupled solves with per-pixel arithmetic using precomputed mask-dependent terms. Reconstructions are parallelized across CPU cores by assigning different compressed frames to different workers per iteration, yielding an ∼15-fold speedup on an Intel Xeon w5-2465X (16 cores) for a representative zebrafish dataset at the standard compression setting (**Supplementary Table 2**). Final reconstructions are saved as 16-bit TIFF stacks.

### Rolling-shutter correction

CAPS measurements acquired with the camera operated in rolling-shutter mode contained line-dependent temporal shear. Because each camera exposure was synchronized to a predefined sequence of DMD masks, this distortion was deterministic in our system. For each compression ratio, we established the correspondence between sensor-line position and mask timing by experimental calibration using shifted DMD illumination patterns (**Supplementary Note 3B**). After each raw frame was reconstructed into CR images, the resulting line-to-time mapping was used to partition the decoded images into ordered line segments and to reassign complementary segments to the physical plane whose acquisition time they most closely matched.

### Fluorescent beads imaging

Fluorescent beads were used for both optical characterization of the synchronized light sheet system and benchmarking of CAPS reconstruction fidelity. For optical characterization, 500 nm fluorescent Nile Red beads (FP-0556-2, Spherotech) embedded in 1% agarose were imaged under 532 nm excitation to measure the illumination profile and PSF during synchronized volumetric scanning. For CAPS fidelity benchmarking, 10–14 µm diameter fluorescent Nile Red beads (FP-10056-2, Spherotech) embedded in 1% agarose were imaged as a static sample using the same CAPS settings as for heart imaging (100 vps over a 190-µm axial range). For each bead dataset, a fully sampled reference volume was acquired over the same axial range by motorized axial scanning at 1 µm steps.

### Synthetic 4D beating phantom generation

A synthetic two-chamber beating phantom was generated as a periodic 4D sequence (50 volumes per cardiac cycle) to emulate zebrafish cardiac dynamics. Each volume was defined on a 400×400×75 voxel grid, with the same lateral pixel size and axial step size as the CAPS acquisitions. The ventricle and atrium were modeled as two prolate ellipsoids connected by a narrow AV junction. Cell nuclei were placed on thin surface shells of each chamber (excluding the overlap region) and rendered as triaxial ellipsoids (5–7 µm diameter) with heterogeneous intensities. Chamber motion was introduced by sinusoidal modulation of atrial and ventricular scale factors in antiphase to mimic reciprocal filling and emptying. The simulated volumes were convolved with a 3D Gaussian kernel to approximate optical blur and corrupted with Poisson noise to mimic photon-limited imaging.

### Zebrafish husbandry and drug treatment

Zebrafish experiments were performed in compliance with protocols approved by the University of Texas at Dallas Institutional Animal Care and Use Committee (IACUC #20–07). Transgenic lines *Tg(myl7:nucGFP)* and *Tg(gata1a:dsRed)* were used to label cardiomyocyte nuclei and erythroid cells, respectively. Embryos were obtained by natural mating and maintained in standard E3 medium, with 0.003% 1-phenyl-2-thiourea (Sigma-Aldrich) added at 1 dpf to suppress pigmentation. Fluorescence-positive larvae were screened at 24 h post-fertilization and imaged at 3 to 4 dpf. Because sex cannot be reliably determined at the larval stages used here, sex was not considered as a variable. Before imaging, larvae were anesthetized in 0.05% tricaine for 10 min and immobilized in 1% low-melt agarose inside a fluorinated ethylene propylene tube, then immersed in a water chamber for imaging; the tube was mounted on rotation stages to optimize heart orientation for CAPS acquisition. For drug treatment experiments, larvae from *Tg(myl7:nucGFP)* or *Tg(gata1a:dsRed)* clutches were randomly assigned at 2 dpf to amiodarone or control groups. Treated larvae were incubated for 24 h in E3 medium supplemented with 0.003% 1-phenyl-2-thiourea and 10 µM amiodarone, whereas control larvae were handled identically in E3 medium with matched vehicle when applicable. Before imaging, larvae were rinsed in fresh E3 and mounted as described above.

### Quiescent and beating zebrafish heart imaging

Quiescent and beating zebrafish hearts were imaged using CAPS. *Tg(myl7:nucGFP)* zebrafish larvae were treated with *tnnt2* morpholino to arrest the heartbeat. The morpholino stock was prepared at 1.0 mM in sterile water, diluted to a 0.5 mM working solution in sterile water containing 0.05% phenol red, and microinjected into embryos at the 1- to 4-cell stage using glass micropipettes. Injected embryos were returned to E3 medium for development and imaged at 3 dpf. CAPS data were acquired at 100 vps over a 190-µm axial range, and a fully sampled reference volume over the same axial range was subsequently acquired by motorized axial scanning at 1 µm steps. For beating heart imaging, transgenic *Tg(myl7:nucGFP)* larvae for cardiomyocyte nuclei and *Tg(gata1a:dsRed)* for erythroid signal were imaged using CAPS. CAPS data were acquired at 100 vps over a 190-µm axial range or 200 vps over a 100-µm axial range. Within each camera exposure, fluorescence was temporally encoded by the DMD using a predefined sequence of binary masks, such that each raw frame contained multiplexed signal from multiple axial positions sampled during that exposure.

### Data preprocessing

Reconstructed stacks were preprocessed to ensure consistent axial sampling and volume indexing (**Supplementary Note 5**). Deviations from the nominal camera timing were first corrected by re-slicing the stack along its frame axis using the measured exposure time, preventing cumulative misassignment of axial positions. Sinusoidal sweeps were then remapped onto a uniform axial grid based on the commanded scan trajectory, and depth-dependent intensity bias arising from nonuniform sweep velocity was normalized. Continuous acquisitions were subsequently split into single-sweep sub-volumes, backward sweeps were reversed to enforce a consistent axial order, and forward/backward volumes were registered to reduce direction-dependent offsets and improve temporal consistency.

### Statistical analysis

Data are presented as mean ± SD unless otherwise noted. When distributions were skewed or when assumptions for mean-based analyses were not met, data are reported as median and IQR (25th–75th percentile), where a smaller IQR indicates lower variability and a larger IQR indicates greater dispersion of the data. Box plots show the median and IQR, with whiskers extending to 1.5 × IQR; individual points represent individual measurements. All statistical tests were two-sided. For comparisons across more than two groups, we used the Friedman test. When the omnibus Friedman test was significant, post hoc pairwise comparisons were performed using paired Wilcoxon signed-rank tests with Holm correction for multiple testing. Statistical significance was defined as P < 0.05.

### Cell segmentation and tracking

Automated 3D instance segmentation and cell tracking of zebrafish cardiomyocytes were performed using BiaPy^42^ and Ultrack^43^. A customized 3D U-Net was trained in BiaPy (feature maps: 32, 64, 128; kernel size: 3; batch normalization; ELU activation; transposed-convolution up-sampling; sigmoid output) using six manually annotated CAPS volumetric datasets. These training datasets were selected to capture biological variability across individual fish and heart morphologies, including different contraction states, thereby improving generalizability. BiaPy was configured to automatically output an optimal segmentation result based on learned probability thresholds. To improve tracking robustness, we extended this workflow by generating segmentation hypotheses for each 4D dataset at three confidence levels: stringent, intermediate and permissive. This provided Ultrack with a broader set of candidate cells and allowed the algorithm to resolve missed detections and over-segmentation errors through temporal consistency. Ultrack then integrated these candidates over time to recover cell tracks while minimizing errors from local under-segmentation, over-segmentation and missed detections. Detailed software configurations and hyperparameter settings for both BiaPy and Ultrack are provided in **Supplementary Table 3**.

### Heart chamber volume quantification

To assign segmented cells to the atrium and ventricle, we partitioned the cardiomyocyte point cloud into two clusters using spectral clustering after cell segmentation^44^. Approximate chamber centers were manually identified across the cardiac cycle to initialize cluster identity and prevent label swapping. Small disconnected outlier clusters were removed, and only the dominant contiguous component was retained to ensure temporal consistency. Chamber volume was estimated from the 3D point cloud of each labeled chamber using a convex hull volume computation^45^, with physical units obtained from the experimental voxel spacing. The resulting atrial and ventricular volume traces were used to quantify cardiac function, including end-diastolic and end-systolic volumes, stroke volume, and ejection fraction.

### 4D blood flow field analysis

To quantify intracardiac blood flow, we analyzed the erythroid channel from *Tg(gata1a:dsRed)* datasets and estimated a flow field directly from the reconstructed volumes using Ultrack^43^. For each pair of consecutive reconstructed volumes, Ultrack computed a displacement vector field by optimizing a multiscale, gradient-based image matching objective that warps the earlier volume toward the subsequent one. Displacements were converted to physical velocities by scaling with voxel dimensions and dividing by the inter-volume time interval, yielding voxel-wise velocity components and speed magnitudes at each time point. For regional quantification, we defined 3D regions of interest in the atrium, AV canal, and ventricle, and extracted mean speed traces within each region of interest across time. Flow directionality was assessed by projecting the velocity vectors onto a user-defined atrium-to-ventricle transport axis, with the sign of the projected component distinguishing antegrade from retrograde transport. Rhythmic structure was characterized by Fourier amplitude spectra of the mean speed traces from each region of interest.

### PSNR, SSIM and NRMSE calculations

Reconstruction quality was quantified using PSNR and SSIM by comparing each reconstruction with the corresponding fully sampled reference. PSNR measures pixel/voxel-wise intensity error and was computed as

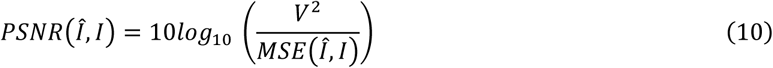

where *Î* and *I* denote the reconstructed and reference images (or volumes), *V* is the maximum intensity value, and MSE is the mean squared error.

SSIM was used to assess preservation of structural information. For a pair of images, *a* and *b*, SSIM was computed as

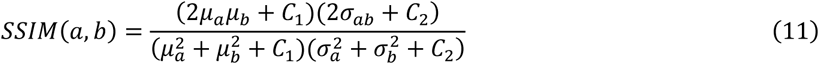

where *µ_a_* and *µ_b_* are local means, *σ_a_ ^2^* and *σ ^2^* are local variances, and *σ_ab_* is the local covariance. Stabilizing constants were set to *C_1_* = (0.01 × *L*)^2^ and *C_2_* = (0.03 × *L*)^2^ with L=1 for normalized images.

For chamber-volume analysis, the agreement between reconstructed chamber volume traces and the reference was quantified using the NRMSE, defined as

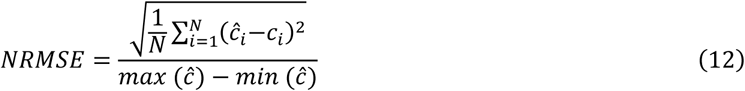

where *ĉ_i_* and *c_i_* denote the reconstructed and reference chamber volumes at phase *i*, respectively, and *N* is the number of matched volumes.

## Data and materials availability

The data and code underlying the results reported in this paper are available from the corresponding authors upon request.

## Acknowledgments

We acknowledge Dr. Caroline Burns’s group at Boston Children’s Hospital, including Drs. Mengmeng Huang and Hui-Min Yin, for generously sharing the transgenic zebrafish. We thank Ms. Elizabeth Ibanez and Mr. Adam Lavitz for their assistance with zebrafish husbandry at UT Dallas. We also appreciate the constructive comments and discussions provided by the D-incubator members at UT Dallas.

## Funding

National Institutes of Health grant R00HL148493 (Y.D.)

National Institutes of Health grant R01HL162635 (Y.D.)

National Science Foundation grant 2503230 (Y.D.)

Cecil H. and Ida Green Professorship in Systems Biology Science (Y.D.)

## Author contributions

X.Z. and Y.D. conceived the idea of CAPS microscopy through coded axial multiplexing and computational reconstruction. X.Z., J.C. and Y.D. designed and built the CAPS hardware. J.C., X.Z., M.A. and Y.D. developed the CAPS control software. X.Z., J.C. and Y.D. performed microscopy experiments. X.Z., Y.G., Y.L. and Y.D. developed the customized PnP-ADMM reconstruction algorithm. X.Z. and Y.D. developed the rolling-shutter correction and data preprocessing pipeline. X.Z., A.S., A.H., K.C. and Y.D. handled zebrafish husbandry and sample preparation. X.Z., J.C. and Y.D. performed cell tracking and flow analysis. X.Z., J.C., Y.G., M.A., J.B., A.S., J.J., K.C., Y.L. and Y.D. validated the experimental and computational results. X.Z., J.C., Y.G., Y.L. and Y.D. carried out the investigation. X.Z., J.C., Y.G., J.B., M.A., A.S., Y.L. and Y.D. wrote the original draft. All authors contributed to review and editing. Y.D. supervised and administered the project. Y.D. acquired funding.

## Supplementary Information

The file includes:

Supplementary Notes 1 to 5

Supplementary Figs. 1 to 13

Supplementary Tables 1 to 3

Legends for Supplementary Videos 1 to 8 References

## Other Supplementary Material for this manuscript includes the following

Supplementary Videos 1 to 8

